# Leak current, even with gigaohm seals, can cause misinterpretation of stem cell-derived cardiomyocyte action potential recordings

**DOI:** 10.1101/2022.10.13.511949

**Authors:** Alexander P. Clark, Michael Clerx, Siyu Wei, Chon Lok Lei, Teun P. de Boer, Gary R. Mirams, David J. Christini, Trine Krogh-Madsen

## Abstract

Human induced pluripotent stem cell-derived cardiomyocytes (iPSC-CMs) have become an essential tool to study arrhythmia mechanisms. Much of the foundational work on these cells, and the computational models built from the resultant data, has overlooked the contribution of seal-leak current on the immature and heterogeneous phenotype that has come to define these cells. Here, we use *in silico* and *in vitro* studies to demonstrate how seal-leak current depolarises action potentials (APs), substantially affecting their morphology, even with seal resistances (R_seal_) above 1GΩ. We show that compensation of this leak current is difficult due to challenges with recording accurate measures of R_seal_ during an experiment. Using simulation, we show that R_seal_ measures: 1) change during an experiment, invalidating the use of pre-rupture values, and 2) are polluted by the presence of transmembrane currents at every voltage. Finally, we posit the background sodium current in baseline iPSC-CM models imitates the effects of seal-leak current and is increased to a level that masks the effects of seal-leak current on iPSC-CMs. Based on these findings, we make three recommendations to improve iPSC-CM AP data acquisition, interpretation, and model-building. Taking these recommendations into account will improve our understanding of iPSC-CM physiology and the descriptive ability of models built from such data.

**Key points:**

- Human induced pluripotent stem cell-derived cardiomyocytes (iPSC-CMs) are an essential tool in the study of cardiac arrhythmia mechanisms.
- Their immature and heterogeneous action potential phenotype complicates the interpretation of experimental data, and has slowed their acceptance in industry and academia.
- We suggest that a leak current caused by an imperfect pipette-membrane seal during single-cell patch-clamp experiments is partly responsible for inducing this phenotype.
- Using *in vitro* experiments and computational modelling, we show that this seal-leak current affects iPSC-CM AP morphology, even under ‘ideal’ experimental conditions.
- Based on these findings, we make recommendations that should be considered when interpreting, analysing and fitting iPSC-CM data.

## 1 Introduction

Human induced pluripotent stem cell-derived cardiomyocytes (iPSC-CMs) are a renewable and cost-effective model for studying cardiac arrhythmia mechanisms in human cells. Patient-specific cells can be used to investigate genetic disease mechanisms (Han et al., 2014), drug cardiotoxicity (Mathur et al., 2015), and inter-patient variability (Blinova et al., 2019). Computational approaches have even been developed to translate experimental results from iPSC-CMs to make predictions in adult cardiomyocytes (Jæger et al., 2021).

Progress in many of these areas, however, has been slowed by the immature phenotype and cell-to-cell heterogeneity of iPSC-CMs (Jonsson et al., 2012; Goversen et al., 2018a). Investigating the source of this variability and its biological implications is important as we come to use iPSC-CMs (and mechanistic models describing their behaviour) for drug safety assessment (Mirams et al., 2016). Recently, Horváth et al. (2018) showed that these limitations can be attributed, at least in part, to the presence of leak current (I_leak_) during patch-clamp experiments. I_leak_ is an experimental artefact caused by an imperfect seal between the electrode pipette tip and cell membrane (Figure 1). Compared to adult cardiomyocytes, typical iPSC-CMs are smaller (leading to a lower membrane capacitance C_m_ < 100pF) and have fewer ion channels. Combined, this makes membrane potential recordings in iPSC-CMs particularly susceptible to imperfect seals. We believe the effects of I_leak_ on the interpretation of iPSC-CM action potential (AP) data has been overlooked by the field, including ourselves (Lei et al., 2017; Clark et al., 2022). Such data have been used in numerous studies to investigate cell-line specific characteristics, and have formed the basis for widely-used iPSC-CM computational models (Paci et al., 2013; Koivumäki et al., 2018; Kernik et al., 2019).

**Figure 1:**
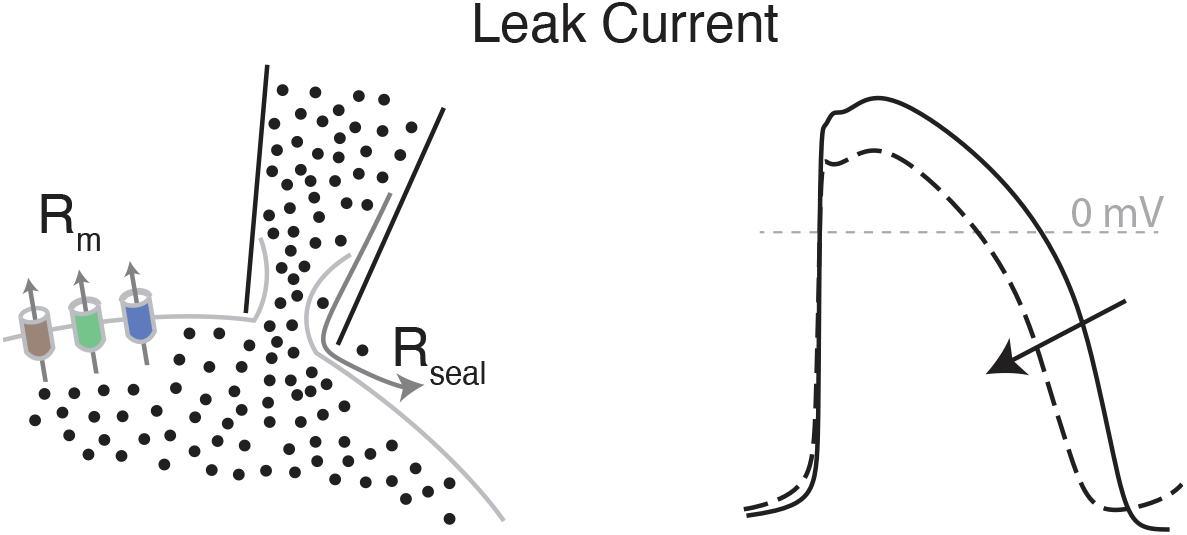
Presence of I_leak_ has undesirable effects on AP morphology and muddles data interpretation. Leak current is an undesirable artefact in patch-clamp experiments. It flows through the seal formed between the pipette tip and cell membrane, and has a magnitude that is inversely proportional to the size of the seal resistance. This artefact affects AP morphology, with greater deviations from baseline (indicated by the arrow) as membrane resistance (R_m_) increases and/or R_seal_ decreases.

In this study, through *in vitro* experiments and computational modelling we show that I_leak_ affects iPSC-CM AP morphology, even under ‘ideal’ experimental conditions. We show that seal resistance (R_seal_) cannot be easily compensated because it cannot be accurately measured during an experiment. Additionally, we posit that the background sodium current (I_bNa_) in iPSC-CM models may be overestimated and mimic the effects of leak on AP morphology. Ultimately, we argue that leak current should be considered when interpreting, analysing, and fitting iPSC-CM AP data.

## 2 Results

### 2.1 Leak affects AP morphology even at seal resistances above 1 GΩ

To investigate the effects of leak current on AP morphology, we added a leak equation to the Kernik (Kernik et al., 2019) and Paci (Paci et al., 2013) models. Knowing that leak acts as a depolarising current in iPSC-CM studies, and lacking information about specific charge carriers, we modelled I_leak_ as having a reversal potential of zero (Ahrens-Nicklas and Christini, 2009; Fabbri et al., 2020):

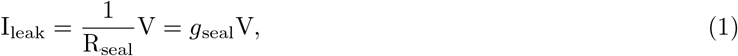

where R_seal_ is the seal resistance and V denotes the membrane potential. The inverse of R_seal_ is a conductance, *g*_seal_, and will be used throughout this study. Note that more complicated equations for leak current (non-linear, and/or with a non-zero reversal potential) may be required in experiments where CaF_2_ seal enhancer is used (Lei et al., 2021).

The effect of I_leak_ on the evolution of V was modelled as:

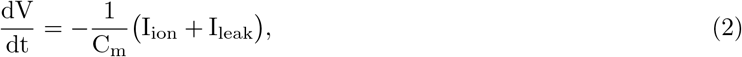

where I_ion_ represents the sum of transmembrane currents and C_m_ is the membrane capacitance.

We used these models to simulate AP recordings with *g*_seal_ set to values between 0.1 nS and 1 nS (i.e., R_seal_ between 10 GΩ and 1 GΩ). The results show that I_leak_ substantially alters AP morphology, even when *g*_seal_ < 1nS, equivalent to R_seal_ ≥ 1GΩ (Figure 2). In simulations with either model, an increase in *g*_seal_ causes depolarisation of the minimum potential (MP), as I_leak_ is inward at negative potentials. Increased leak also causes a substantial reduction in the maximum upstroke velocity, dV/dt_max_, likely due to an incomplete recovery of sodium channels at these depolarised MPs. Interestingly, I_leak_ effects on the action potential duration at 90% repolarisation (APD_90_) differ for the two models — increased *g*_seal_ causes AP shortening in the Kernik model and AP prolongation in the Paci model. These differences are likely due to differences in the relative size of I_leak_ compared to the other repolarising currents during phases one and two of the AP. There are also differences in the effect of *g*_seal_ on cycle length (CL): In the Kernik model, increases in *g*_seal_ lead to a decrease in CL. The Paci model shows more complex dynamics — increases in *g*_seal_ initially lead to prolongation, but then shortening as *g*_seal_ approaches 1 nS.

**Figure 2:**
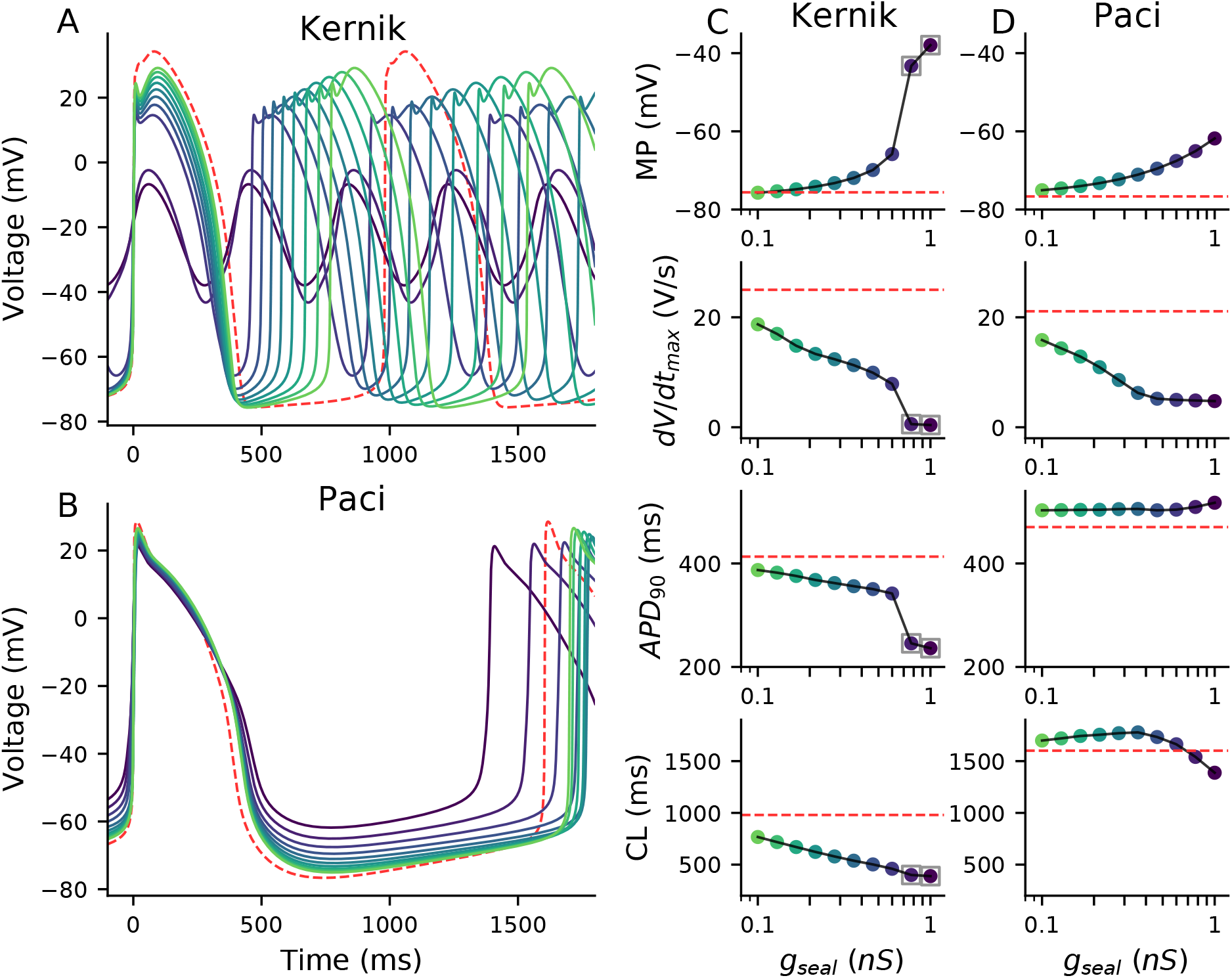
Effect of seal on Kernik and Paci APs. Simulations from the Kernik+leak (**A**) and Paci+leak (**B**) models, each with capacitance set to 98.7 pF (i.e., Paci baseline value), and *g*_seal_ set to values from 0.1 nS to 1 nS (i.e., from 10 GΩ down to 1 GΩ). The dashed red trace shows a baseline (leak-free) simulation. Four AP morphology metrics for the Kernik (**C**) and Paci (**D**) models are plotted against *g*_seal_ (displayed on log-scaled x-axis): minimum potential (MP), maximum upstroke velocity (dV/dt_max_), action potential duration at 90% repolarisation (APD_90_), and cycle length (CL). Grey boxes denote the metrics from the two Kernik simulations that did not produce full APs.

Given these model predictions, it appears likely R_seal_ can alter AP morphology, even at values above 1 GΩ (i.e., below 1 nS). This finding points to the importance of recording accurate measures of R_seal_, and the need for a strategy to address I_leak_ effects during experiments. In the following sections, using *in silico* and *in vitro* data, we show the challenges of devising such a strategy and how, under certain conditions, it may be impossible to determine R_seal_.

### 2.2 R_seal_ is not stable

Unlike voltage-clamp recordings, the effects of I_leak_ on AP morphology (measured in current clamp mode) cannot be corrected in post-processing. Current-clamp leak compensation requires the real-time injection of a current that opposes I_leak_ at every instant during an action potential. This can be achieved using dynamic clamp, but requires a reliable approximation of R_seal_ to recover the leak-free phenotype.

R_seal_ is typically estimated before gaining access to a cell. It can be calculated using a small test pulse in voltage-clamp mode (HEKA Elektronik GmbH, 2016):

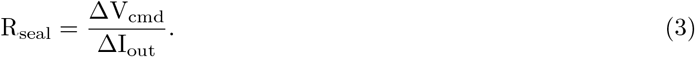

Here, ΔV_cmd_ is the applied voltage step and ΔI_out_ is the difference in recorded current before and during the step. Once access is gained to a cell it can be difficult to estimate R_seal_, as the measured input resistance (R_in_) depends on both R_m_ (membrane resistance) and R_seal_ (Equation 4, Figure 3):

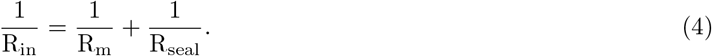

Since R_seal_ is difficult to determine during experiments with iPSC-CMs, it is tempting to measure the value before gaining access and assume it remains unchanged for the duration of an experiment. To investigate this, we considered *in vitro* R_in_ measures taken at two times during iPSC-CM experiments. Rin was measured with 5mV steps from a holding potential of 0mV (i.e., the leak reversal potential) before and after acquiring current clamp data. Due to the large range of R_in_ measures (0.182 GΩ to 52 GΩ), we chose to display the distribution of this data as 1/R_in_, or *g*_in_ (Figure 4A). The data are skewed, with a mean of 1.43 nS (R_in_=0.70 GΩ) and median of 1.19 nS (R_in_=0.84 GΩ).

**Figure 3:**
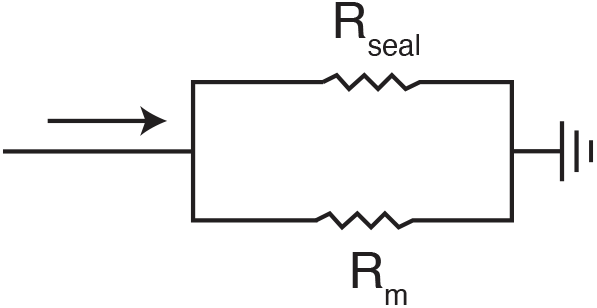
R_seal_ cannot be measured directly once access is gained. Once access is gained, we can only measure the combined resistance R_in_, which is equal to the parallel resistances of R_seal_ and R_m_ (Equation 4). The presence of Rm introduces uncertainty when Rin is used to approximate Rseal, making it difficult to accurately correct for leak current effects. For simplicity, we have omitted other elements of this patch-clamp diagram (e.g., series resistance, capacitance, etc.).

**Figure 4:**
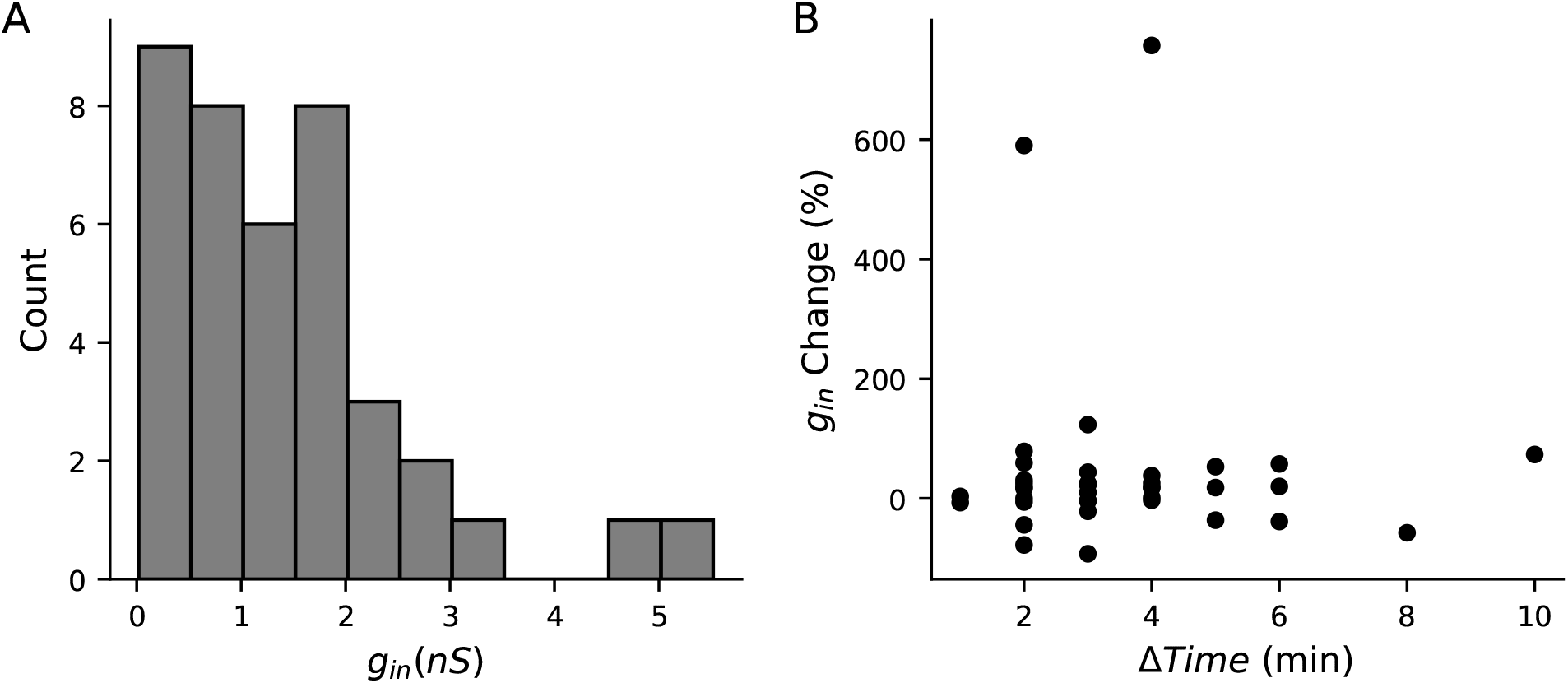
R_in_ changes during iPSC-CM experiments. **A**, Distribution of initial *g*_in_ measurements from iPSC-CMs acquired with a +5mV step from 0mV (n=39). **B**, The percentage change in *g*_in_ plotted against the time elapsed between *g*_in_ measurements. Data is shown for all cells where two *g*_in_ measures were available (n=38). The interval between measurements ranged from 1 to 10 minutes. Time was recorded to the nearest minute, leading to the appearance of banding in the Δ*Time* measure.

The relative change in *g*_in_ from the first to the second time point was calculated, and is plotted against the time elapsed between *g*_in_ measurements in Figure 4B. The mean increase in *g*_in_ was +46% (with a standard deviation of 155%) and the median increase was +18%. Because positive and negative changes cancel each other out in these statistics, we also inspected the absolute change, where we found a mean of 67% (a standard deviation of 147%) and a median of 25%.

The data in Figure 4B illustrate that Rin measurements often change over time. If we assume Rm is stable during experiments, this change in R_in_ should be be attributed to R_seal_, and suggests that the average cell’s I_leak_ increases over time. These findings demonstrate that pre-rupture Rseal measures cannot be taken as ground truth throughout an experiment. As a result, it becomes desirable to find accurate measures of Rseal and Rm after access is gained.

### 2.3 R_in_ is not a good approximation of R_seal_ at any holding potential

In Figure 4 we showed R_in_ measurements from a holding potential of 0 mV. A holding potential of −80mV is a common choice for approximating R_seal_ with R_in_ measures. At this potential, sodium, calcium, and several potassium currents are expected to be largely inactive, but contributions from both I_K1_ and I_f_ must still be considered.

We recently showed that I_f_ is present in at least some of the iPSC-CMs used in this study (Clark et al., 2022). I_f_ is also present in both the Kernik and Paci models, and we found the dynamics of the Kernik I_f_ model to be quite similar to the *in vitro* data in this study (Figure 5A-B). Figure 5A shows a typical cell’s response to an If-activating hyperpolarising step before and after treatment with quinine, at a concentration expected to lead to 32% I_f_ block (this data is taken from a section of a larger protocol — see Figure 6A of Clark et al., 2022). A change in total current of 2 A/F is observed after holding near −120mV for 1 s (Figure 5A). Simulations using the Kernik model with 32% block of I_f_ show a similar directional change, but only a 1 A/F shift in I_ion_ (Figure 5B). Given that most currents, besides I_K1_ and If, are not active at −120 mV, and quinine does not block I_K1_ at the concentration used in the study, we assume the 2 A/F change is due entirely to I_f_ block. Following from this assumption, we can say the cell’s I_f_ conductance per unit capacitance is approximately twice as large as in the baseline Kernik model.

**Figure 5:**
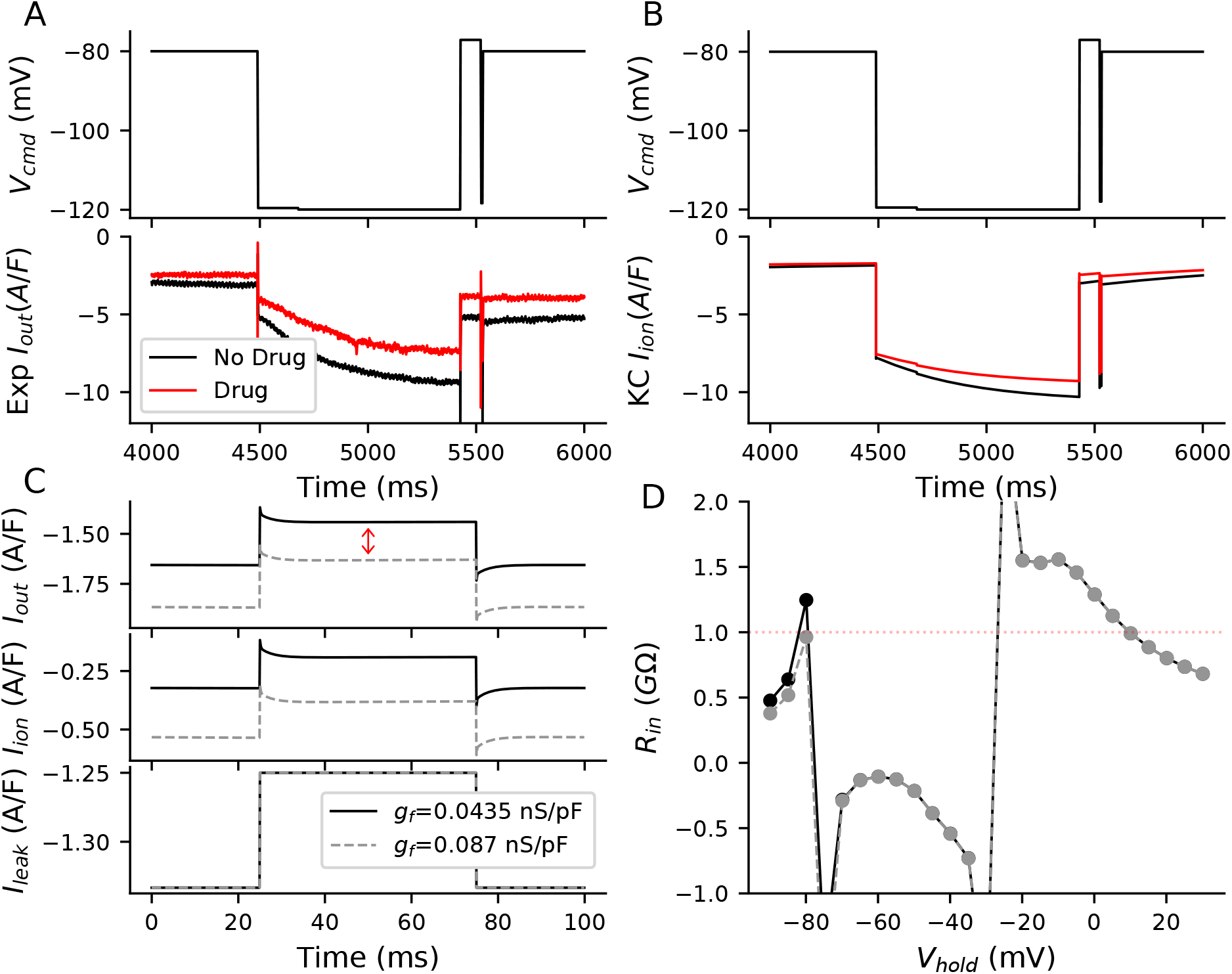
Ignoring the presence of I_f_ makes it impossible to accurately measure R_seal_ after gaining access. **A**, Voltage clamp data acquired from an iPSC-CM before and after treatment with quinine, which is expected to block 32% of I_f_ at the concentration used. **B**, Kernik model response at baseline and with 32% block of I_f_. **C**, Kernik+leak voltage clamp simulations conducted with R_seal_=1 GΩ, *g*_K1_ reduced by 90%, and *g*_f_ set to 0.0435 nS/pF (solid line) or 0.087 nS/pF (dashed line). A voltage step from −80 mV to −75mV was applied, as is commonly used to estimate Rin. This Rin value is sometimes used to approximate Rseal when the holding potential is near −80mV. The amplifier-measured (I_out_), total transmembrane (I_ion_), and leak currents (I_leak_) are displayed. The red arrow (top) indicates the change in I_out_ caused by the different *g*_f_. The R_in_ values calculated based on ΔI_out_ are 1.25 GΩ and 0.96 GΩ for the 0.0435 nS/pF and 0.087 nS/pF simulations, respectively. **D**, R_in_ values are plotted against holding potential for Kernik+leak models with R_seal_ = 1 *G*Ω and *g*_f_=0.0435 nS/pF or *g*_f_=0.087 nS/pF. The red dotted line shows the true simulated R_seal_ value of 1 GΩ.

**Figure 6:**
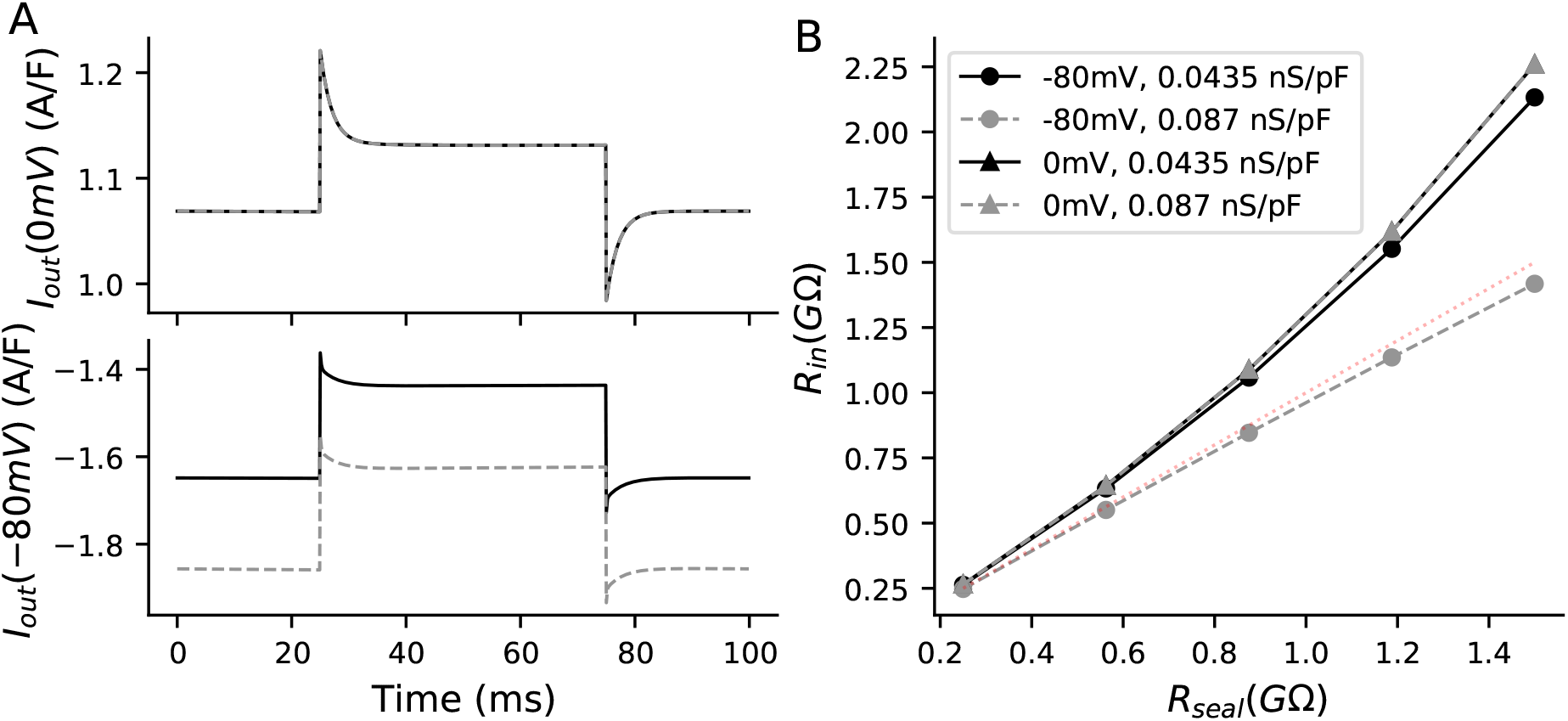
R_in_ predictions of R_seal_ are overestimated at the reversal potential for leak current. **A**, The current response (I_out_) for Kernik+leak models with a 1GΩ seal and *g*_f_ of 0.0435 nS/pF (solid line) or 0.087 nS/pF (dashed line) to a 50 ms +5 mV voltage clamp step from 0mV (top) or −80mV (bottom). **B**, Effect of R_seal_ on R_in_ measures for models with *g*_f_ set to 0.0435 (solid) or 0.087 (dashed) nS/pF. R_in_ was calculated with Equation 3. The +5 mV voltage steps were taken from either 0 or −80 mV. The R_seal_ = *R*_in_ line (red dotted) is provided as a reference for when R_in_ correctly predicts R_seal_. The 0mV lines are overlapping, illustrating that R_in_ is not sensitive to *g*_f_ at this voltage.

To illustrate the effect of I_f_ on leak calculations, we compared simulations from Kernik+leak models with R_seal_ = 1 GΩ and with *g*_f_ set to either the Kernik baseline value (*g*_f_ = 0.0435 nS/pF) or twice its baseline value (*g*_f_ = 0.087 nS/pF) (Figure 5C). To highlight that I_f_ effects on R_in_ are largely independent of I_K1_, we also reduced the *g*_K1_ in these models to 10% of its original value. The calculated R_in_ values for these models at −80 mV are 1.25 GΩ for *g*_f_=0.0435 nS/pF (baseline) and 0.96 GΩ for *g*_f_=0.087 nS/pF (2 × baseline) — in other words a +25% and −4% error in R_seal_ prediction (Figure 5C). These simulations show that, at −80 mV: 1) I_f_ contributes to I_out_ and affects measures of I_leak_, and 2) R_seal_ may be over- or under-estimated depending on the value of *g*_f_.

We then calculated R_in_ values at multiple holding potentials between −90 and +30 mV to determine whether we could find a potential where R_in_ is close to R_seal_, thereby minimising the prediction error (Figure 5D). The model predicts that 10 mV (R_in_ =0.99 GΩ) minimises the error in our approximation of R_seal_. This makes intuitive sense, as these potentials overlap with the large R_m_ plateau phase of the AP. This does not however, mean that Rin measurements at 10 mV will always produce the best estimate of Rseal. Instead, it indicates the size of Iion does not change much when taking a 5 mV step from this potential. There is, however, a considerable amount of total current present, making this Rseal prediction sensitive to variations in the predominant ionic currents at this potential. Moreover, I_leak_ will be small and therefore more difficult to measure as 10 mV is close to the leak reversal potential (0 mV). It is also worth noting that the complex voltage- and time-dependent behaviour of transmembrane currents make Rin measures sensitive to both the duration and size of the voltage step (see e.g. the supplement to Clerx et al., 2021). In summary, it is difficult to find a holding potential where Rseal can be measured without contamination from any transmembrane currents (i.e., where I_leak_ = I_out_).

Taken together, these findings provide evidence to the claim that Rseal cannot be reliably measured in iPSC-CMs once access is gained.

Next, we compared the effect of I_f_ on R_m_, and the error in assuming R_seal_ ≈ R_in_, at both a 0mV and −80 mV holding potential. At 0 mV the Kernik+leak model is not sensitive to changes in *g*_f_, as I_f_ is largely non-conductive (Figure 6A). However, due to an increased relative contribution of inward currents at 0 mV, the Kernik+leak model predicts an R_in_ with a large overestimation of R_seal_ (Figure 6B). This error increases as the true value of R_seal_ increases. Figure 6B also illustrates the sensitivity of the model to variations in *g*_f_ at −80 mV, with the 0.087 nS/pF model producing a small underestimate of R_seal_ while the 0.0435 nS/pF model overestimates R_seal_ these errors increase as R_seal_ increases. The improved prediction accuracy of the 0.087 nS/pF model at −80 mV is a coincidental side-effect of doubling *g*_f_: with a different distribution of ion current densities or a larger baseline gf value, the same doubling could just as easily worsen R_seal_ predictions. For example, the R_in_ of an iPSC-CM with a large I_K1_ current may slightly underestimate R_seal_ at −80 mV — doubling *g*_f_ in this case would result in a greater underestimation, increasing the error of the estimate.

### 2.4 C_m_ and R_in_(0mV) correlate with minimum potential

The iPSC-CMs used in this study displayed a heterogeneous phenotype (Figure 7), producing both spontaneously firing (n=27) and non-firing (n=12) current clamp recordings. Figure 7A shows three cells with very different baseline current-clamp recordings: non-firing and depolarised (grey), spontaneously firing with a short AP (black), and spontaneously firing with a long AP (blue). Non-firing cells (MP = −42±8 mV) and cells with spontaneously-firing APs were depolarised (MP = −55 ± 7 mV) — the spontaneously-firing cells also had a shorter AP duration (APD_90_ = 133±73 ms) (Figure 7B) relative to adult cardiomyocytes (O’Hara et al., 2011) and iPSC-CM models (Kernik et al., 2019; Paci et al., 2013).

**Figure 7:**
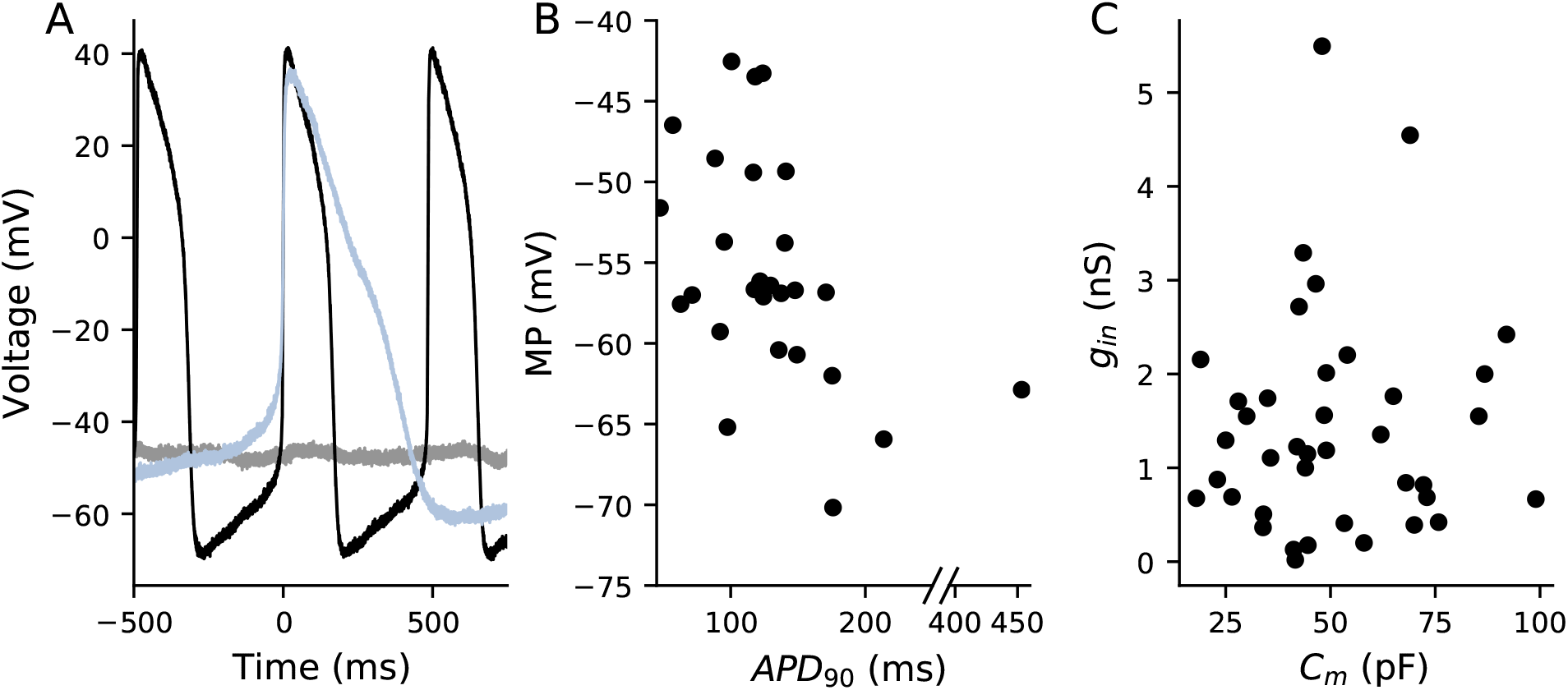
Cells appeared phenotypically heterogeneous, with uncorrelated variation in *g*_in_ and C_m_. **A**, Current clamp recordings from three cells show phenotypic heterogeneity: non-spontaneous (grey), spontaneous AP with short APD (black), and spontaneous AP with long APD (blue). **B**, MP and APD_90_ for spontaneously beating cells (n=27). Note the broken *x*-axis which just allows us to display an outlying data point. **C**, The relationship between C_m_ and *g*_in_ for all cells (n=39).

In this section, we use linear regression analyses to determine if there is a correlation between g_in_/C_m_ and AP biomarkers. The values of each cell’s g_in_ and C_m_ are shown in Figure 7C. I_leak_’s effect on AP morphology is expected to scale directly with g_in_ and inversely with C_m_. This is because *g*_in_, even if a poor estimate, is expected to correlate with *g*_seal_ (Figure 6B). Additionally, a given *g*_leak_ will cause a smaller contribution in larger cells (i.e., cells with larger C_m_), because the ionic currents are expected to scale with the size of the cell.

Four AP biomarkers (MP, APD_90_, CL, and dV/dt_max_) were compared to g_in_/C_m_ (Figure 8). The MPs of spontaneously firing (R=0.47, p<.05) and non-firing (R=0.76, p<.05) cells are positively correlated with *g*_in_/C_m_ (Figure 8A). This finding is in agreement with our *in silico* studies showing that increasing *g*_in_ will depolarise the cell (Figure 2). The other three biomarkers failed at least one of the assumptions that is required when conducting a linear regression analysis (see Methods). There are no obvious trends when comparing *g*_in_/C_m_ to CL or dV/dt_max_. The APD_90_ plot, however, indicates there may be some AP shortening as *g*_in_/C_m_ increases. Due to undersampling and a lack of linearity, we cannot make any claims of significance between these two measures. Leak simulations with the models, though correlated, did not predict a linear relationship between *g*_seal_ and these biomarkers (Figure 2C-D). However, the MP vs. *g*_in_/C_m_ relationship passes all tests of linear regression assumptions and trends in the same direction as the Kernik and Paci simulations in Figure 2.

**Figure 8:**
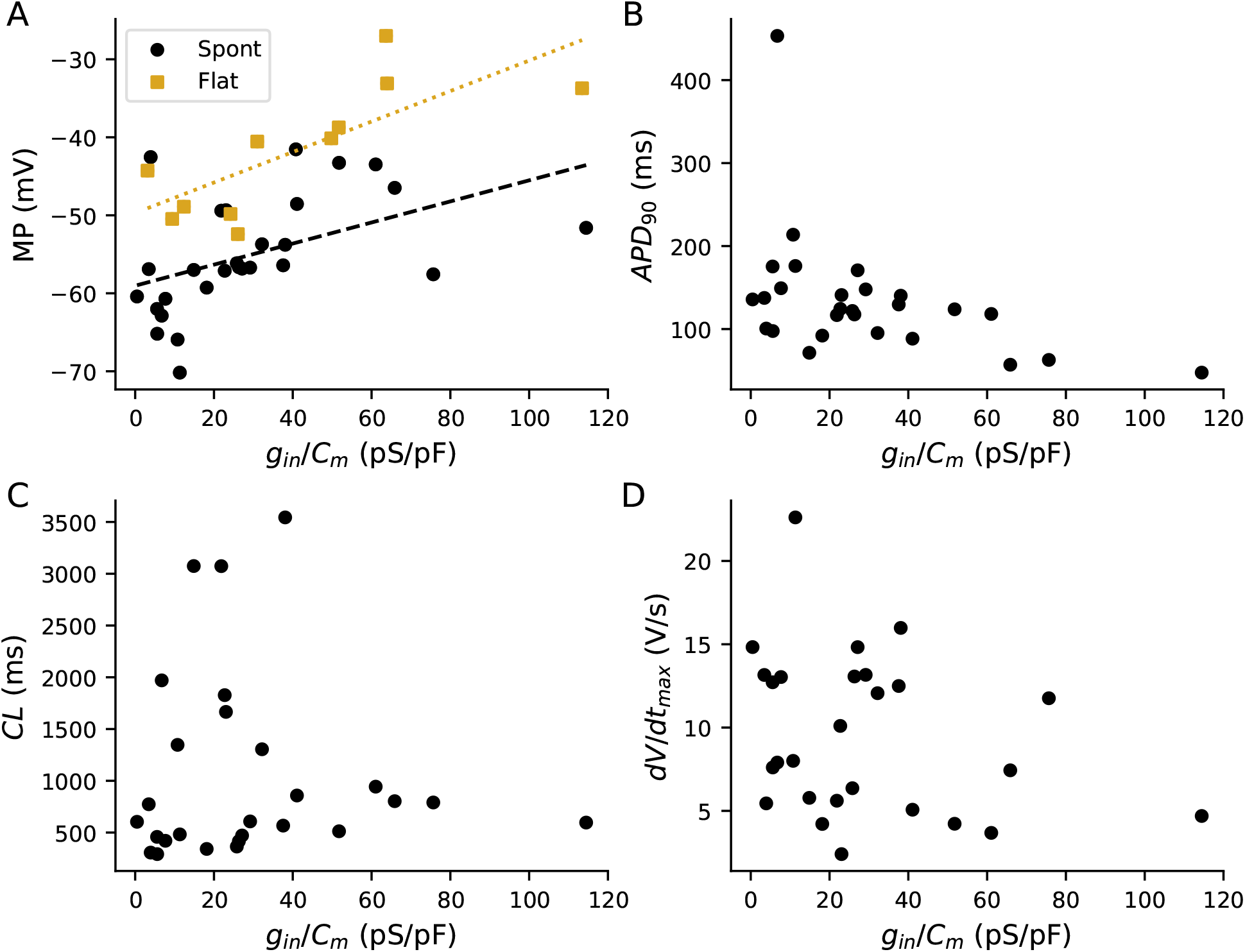
Relationship between *g*_in_/C_m_ and AP biomarkers. **A**, *g*_in_/C_m_ plotted against MP. Spontaneously firing cells are denoted as black points and non-firing cells as yellow squares. Linear regression fits to data from spontaneous (black dashed, *R* = 0.47, *p* < 0.05) and non-firing (yellow dotted, *R* = 0.76, *p* < 0.05) cells are overlaid on the plot. No statistically significant relationship was found between *g*_in_/C_m_ and APD_90_ (**B**), CL (C), or dV/dtmax (**D**).

In summary, we found that a higher *g*_in_/C_m_ (indicating greater I_leak_ contribution) correlated with more depolarised MPs. This supports the idea that I_leak_ affects AP shape and cell-to-cell variability in the iPSC-CMs used in this study.

### 2.5 Fitting background currents in iPSC-CM models can absorb and imitate I_leak_

iPSC-CM models contain linear background currents (sodium and calcium) that differ from I_leak_ only in terms of reversal potentials and the ions they conduct. However, their presence and their magnitudes have not been experimentally investigated in iPSC-CMs (see Discussion). Here, we show that the background currents in existing iPSC-CM models can imitate I_leak_, and we hypothesise that the contribution of I_leak_ may erroneously be ascribed to background currents when models are fit to experimental data.

We used a genetic algorithm (GA) to study the potential of background currents to imitate leak effects (see Methods). We fit the baseline Kernik model to a Kernik+leak model with R_seal_ = 5 GΩ (Figure 9), allowing only the background sodium (*g*_bNa_) and background calcium (*g*_bCa_) conductances to vary. These currents were selected because they were incorporated into the Kernik model without independent iPSC-CM experimentation or validation. The best fit individual had an increased *g*_bNa_ (×7.0), while *g*_bCa_ (×1.0) did not change much relative to the baseline model (Figure 9A). While not a perfect match, the best-fit trace reproduced qualitative features of the baseline+leak trace, showing a depolarised MP and a smaller amplitude (Figure 9B). This indicates that increased I_bNa_ can affect the AP in a fashion similar to I_leak_.

**Figure 9:**
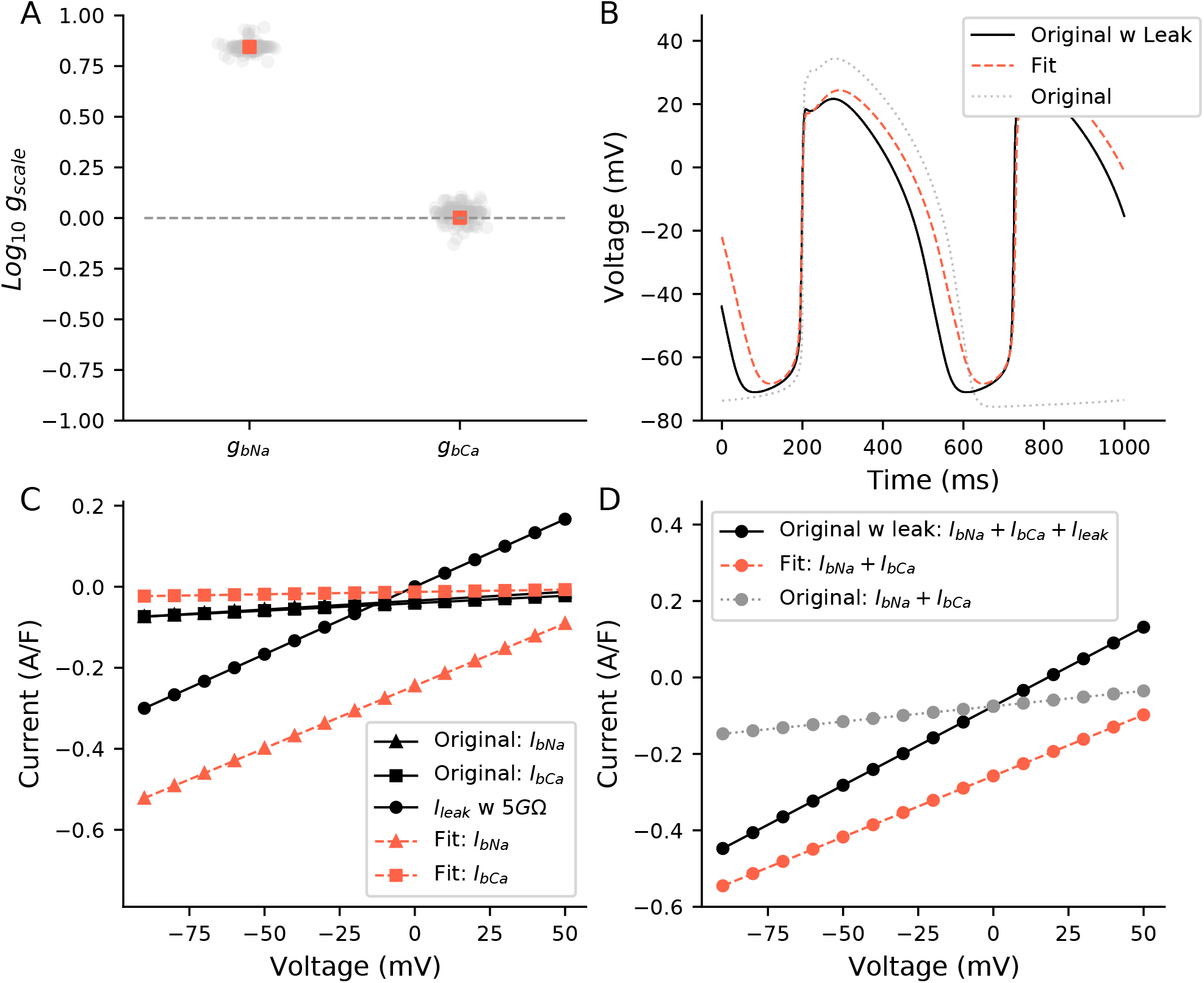
A simulated example of how leak can be absorbed into background currents: Kernik baseline model fit to Kernik+leak model. The I_bNa_ and I_bCa_ conductances (*g*_bNa_, *g*_bCa_) of the baseline Kernik model were fit to a Kernik+leak model (i.e., original+leak) with R_seal_ set to 5 GΩ. A GA with a population size of 150 individuals and 20 generations was used to fit the model. **A**, The conductances for all individuals (grey) and the best fit individual (red square) from the last generation. **B**, Traces from the original baseline Kernik+leak model with a 5 GΩ seal (black), the best fit model from the last generation (red dashed), and the original baseline Kernik model (grey dotted). **C**, IV curves for baseline *I*_bNa_, *I*_bCa_, and I_leak_, and for fitted *I*_bNa_ and *I*_bCa_. **D**, IV curves for: 1) *I*_bNa_ + *I*_bCa_ + *I*_leak_ from the original+leak model, 2) Fitted *I*_bNa_ + *I*_bCa_ from the fitted Kernik model, and 3) *I*_bNa_ + *I*_bCa_ from the original baseline Kernik model.

We then compared the background current IV curves of the fit model to the original baseline+leak model (Figure 9C-D). The IV curves of I_bNa_ (E_Na_=+79 mV) and I_bCa_(E_Ca_=+112 mV) are negative at all tested voltages (−90 to +60 mV), while I_leak_ reverses at 0mV (Figure 9C). The best-fit model I_bNa_ conducts a much larger negative (i.e., depolarising) current at all tested voltages when compared to I_leak_.

We also investigated composite IV curves for: 1) *I*_bNa_ + *I*_bCa_ + *I*_leak_ from the original baseline+leak model, 2) fitted *I*_bNa_ + *I*_bCa_ from the best fit model, and 3) *I*_bNa_ + *I*_bCa_ from the original baseline model (Figure 9D). The fitted background IV curve (red) is negatively shifted, relative to the original baseline model (grey), as the *I*_bNa_ component conducts large depolarising currents, mimicking the effects of I_leak_ at large negative potentials. Despite good AP agreement (Figure 9B), the divergence in the Kernik+leak and fitted model IV curves illustrates that the depolarising effects of I_leak_ and I_bNa_ at negative voltages are most likely responsible for the morphological agreement seen throughout the cycle length (Figure 9D).

In this section, we have shown that I_bNa_ can affect AP morphology in a similar way to I_leak_. Here, the Kernik model was used as our ground truth, but was constructed using iPSC-CM data that may have been polluted by leak current. Unless I_leak_ is explicitly handled, either by experimental real-time dynamic clamp leak correction or in the mathematical model itself at the time of its construction, mathematical iPSC-CM models may absorb the effects of I_leak_ by erroneously increasing background currents.

## 3 Discussion

Leak current is a common and unavoidable experimental artefact that affects patch-clamp recordings. In this study, using both model predictions and experimental data, we show that leak current: 1) affects iPSC-CM AP morphology; 2) can vary during experiments; 3) cannot be accurately estimated after access is gained to an iPSC-CM; and 4) may be absorbed by linear equations for background currents when iPSC-CM models are fit to experimental AP data. During iPSC-CM current-clamp studies, leak consideration often starts with a pre-rupture seal measurement (with a 1 GΩ threshold) and is ignored if the seal appears to remain stable throughout the study. Here, we argue leak effects should be quantitatively scrutinised at all points during the acquisition, analysis and fitting of experimental data. Furthermore, we believe cell-to-cell variation in seal resistance contributes to observed iPSC-CM AP heterogeneity — often attributed nearly entirely to variations in ionic current densities.

### 3.1 Leak affects AP morphology

Leak is known to affect the shape of AP morphology in small cardiac cells. Simulations in chick embryonic cells (with model C_m_ = 25.5 pF) have shown that leak current will substantially depolarise the MP and shorten the CL, even with large R_seal_ values (5 GΩ, Krogh-Madsen et al., 2005). More recently, Horváth et al. (2018) showed that *in vitro* iPSC-CMs were depolarised during single-cell experiments, but not when cells were clustered. These results indicate that iPSC-CMs are not inherently depolarised, but may be affected by the increased influence of leak current in isolated cells with a low capacitance.

Our *in vitro* and *in silico* findings support this conclusion and strengthen the argument that iPSC-CM AP morphology is strongly affected by leak current.

iPSC-CMs have long been defined by their immature and heterogeneous phenotype (Ma et al., 2011; Doss et al., 2012). Over the years, optimisations of the differentiation and dissociation processes have improved cell maturity and consistency, but issues remain (Herron et al., 2016). Such shortcomings of the cells have often been attributed to variations in ionic current conductances and a reduced I_K1_ density (Ma et al., 2011; Doss et al., 2012). However, even iPSC-CMs with large I_K1_ have displayed depolarised MPs and large cell-to-cell variability (Horváth et al., 2018; Feyen et al., 2020). In addition to ionic current densities, we suggest that variations in leak current play a critical role in both the heterogeneity and apparent immaturity (characterised by depolarised MPs) of these cells.

### 3.2 Predicting R_seal_ during experiments

Useful implementation of a leak compensation current requires accurate measures of R_seal_ throughout an experiment. R_seal_ can be well-approximated prior to gaining access, but after perforation (or rupture) the presence of membrane currents make it impossible to obtain an accurate measurement (Figure 5). This is problematic due to the tendency of R_seal_ to change over the course of an experiment (Figure 4).

To address these difficulties, we believe it may be feasible to use the pre-rupture R_seal_ and post-rupture R_in_ measures to calculate estimates of R_seal_ during an experiment. This approach would require an accurate measure of R_in_ just after access is gained. Using R_seal_ and the initial R_in_, it is possible to calculate R_m_ (Figure 3). An estimate of R_seal_ could then be made at any time during the experiment, assuming the calculated R_m_ stays constant, by re-measuring R_in_ and using Equation 4. This approach relies on two major assumptions: 1) the perforation/rupture step does not affect the seal; and 2) a protocol or procedure exists that can be used prior to each measurement of R_in_ to ensure that the contribution of R_m_ is consistent. We cannot say for certain that these assumptions will always be valid. However, we believe that recording frequent R_in_ measurements, estimating R_seal_, and scrutinising changes are important steps for the correct interpretation of iPSC-CM current clamp data.

### 3.3 Correcting for R_seal_ during experiments

We believe these Rseal estimates should be used in a dynamic clamp leak compensation setup to address the limitations caused by a depolarised and variable MP. The approach works by injecting simulated currents into a cell in a real-time continuous loop during current clamp experiments (Ortega et al., 2018). I_K1_ dynamic clamp has been used on iPSC-CMs to attain quiescence at a MP below −70mV so the cells can be paced at a desired frequency (Meijer van Putten et al., 2015; Goversen et al., 2018b; Li et al., 2021; Clark et al., 2022). A dynamically clamped leak-compensation current has also been implemented and used in manual patch-clamp studies with neonatal mouse cardiomyocytes (Ahrens-Nicklas and Christini, 2009), demonstrating the potential of using such an approach with small cardiomyocytes. The effects of leak and the ability of leak compensation to recover adult cardiomyocyte behaviour has also been demonstrated in an *in silico* study (Fabbri et al., 2020). Together, these investigations demonstrate the potential of dynamic clamp as an experimental tool to simultaneously address shortcomings of the cells (i.e., I_K1_ density) and experimental setup (i.e., I_leak_). This technique has the potential to improve the descriptive ability of iPSC-CMs when used in biophysical and drug investigations.

Inaccuracies in these estimates, however, will remain, resulting in the potential to under- or overcompensate. Undercompensation, while an improvement over no compensation, will still result in a depolarised MP and shortened AP duration relative to its true value. Overcompensation will hyperpolarise the MP relative to its true value, but also prolong phases 1 and 2 of the AP. This is because leak compensation is an inward current at positive voltages. Due to the prolongation caused by overcompensation, we believe undercompensation is preferable. We suggest injecting a fraction of the full compensatory current to mitigate the risk of underestimating Rseal. The Nanion Dynamite^8^ sets the leak percent compensation to 70%, which seems reasonable (Becker et al., 2020). Further investigation is needed to provide advice on how to choose this value in all circumstances.

### 3.4 Background currents absorb leak effects

Ion-specific background currents in the Kernik and Paci iPSC-CM models were taken from the ten Tusscher et al. (2004) model. These currents can trace their roots to the seminal work of Luo and Rudy (1994). The currents were included in the Luo and Rudy (1994) and ten Tusscher et al. (2004) models to help to maintain physiologically realistic intracellular concentrations. In the ten Tusscher et al. (2004) model, these currents helped to produce [Na^+^]_i_ frequency changes in line with *in silico* cardiac simulations (Boyett and Fedida, 1988), and equilibrium concentrations within the ranges from an *in vitro* study with human cardiomyocytes (Pieske et al., 2002).

Direct measurements of I_bCa_ and I_bNa_ in iPSC-CMs have not been reported. The Kernik and Paci iPSC-CM models both adopted the ventricular ten Tusscher et al. (2004) formulation for I_bCa_ and I_bNa_, and then set the conductances of these currents by comparing model predictions of the AP with in *in vitro* measurements in iPSC-CMs. We posit that I_bNa_ is overestimated and compensates for the absence of leak current, a source of discrepancy between these models and reality. We expect inclusion of leak when constructing iPSC-CM models to reduce background sodium and result in a more realistic model of *in vitro* single-cell iPSC-CMs.

### 3.5 Modelling experimental artefacts

While the effects of experimental artefacts in single-cell studies are well-established, consideration of them while building ion channel and action potential models has been limited (Whittaker et al., 2020). *In silico* studies investigating series resistance effects on voltage clamp recordings have been done in fast-activating currents, such as I_Na_ and I_to_ (Ebihara and Johnson, 1980; Montnach et al., 2021), but to our knowledge artefact equations have not been included in the calibration process for widely-used models of these currents — although the I_Na_ model by Ebihara and Johnson was incorporated directly into the widely copied INa model by Luo and Rudy (1994). Recently, Lei et al. (2020) demonstrated that coupling experimental artefact equations with an I_Kr_ mechanistic model improved predictions. These studies show that experimental artefact equations can improve the descriptive ability of electrophysiological models. As such, we believe experimental artefacts should be explicitly taken into account at the modelling phase, and not ignored simply because a pre-determined minimum threshold is reached (e.g., 1 GΩ). Based on the findings of our study, we believe cardiomyocyte models, and especially iPSC-CM models, should explicitly include leak currents when fitting to experimental current clamp data.

### 3.6 Recommendations

The results in this manuscript provide important insights for experimentalists and modellers alike. We developed the following recommendations based on our findings:

1. *Experimental:* R_seal_ should be recorded before gaining access to a cell, and R_in_ should be measured frequently during an experiment. It is important to measure R_in_ from a voltage that provides a consistent measure of R_m_, such that any changes in R_in_ can be attributed to changes in R_seal_. If these measures of R_in_ do not vary, this may be indicative of a stable R_seal_.
2. *Experimental:* Dynamic injection of a leak compensation current can help the cell recover its native MP and produce an AP with little contribution from I_leak_. Because R_seal_ is difficult to measure during experiments, and to avoid overcompensation, we advise injecting a current that compensates for a fraction (e.g., 70%) of the estimated I_leak_. Additionally, the R_seal_ and R_in_ measures should be reported along with iPSC-CM data.
3. *Modelling:* Inclusion of the I_leak_ equation will improve the descriptive ability of iPSC-CM models. While this equation may not always improve fits to AP data, it will take into account an important current affecting iPSC-CM recordings.

### 3.7 Limitations and future directions

This study has several limitations that should be considered during future investigations that may be affected by I_leak_. First and foremost, when gathering these data for a previous study we did not follow our own recommendation of recording the exact value of R_seal_ before gaining access and then measuring R_in_ just after perforation. In the future, we hope to use these two values to predict R_seal_ at multiple timepoints during an experiment, as outlined in Section 3.2. Second, we only conducted these experiments in one cell line. While our results appear similar to data from other labs (e.g., Horváth et al., 2018), it would be useful to conduct this study on multiple cell lines in the same lab. Third, we did not attempt dynamic injection of a leak compensation current — in future work we would like to investigate this as an approach to reducing cell-to-cell heterogeneity. Finally, the iPSC-CM models have innumerable differences from the cells used in this study, which is evident when comparing AP morpoholgies of *in vitro* cells (Figure 7A) to *in silico* models (Figure 2). However, agreement that we did see between simulations and our *in vitro* data demonstrate the potential of improving the descriptive ability of iPSC-CM models by including a leak current.

### 3.8 Conclusion

In this study, we demonstrate that leak current affects iPSC-CM AP morphology, even at seal resistances above 1 GΩ, and contributes to the heterogeneity that characterises these cells. Using both *in vitro* and *in silico* data, we showed the challenges of estimating R_seal_ after gaining access to a cell and that R_seal_ is subject to change during the course of an experiment. We also posit that background sodium current in iPSC-CM models may be responsible for masking leak effects in *in vitro* data. Based on these results, we made three recommendations that should be considered by anyone who collects, analyses, or fits iPSC-CM AP data.

## 4 Methods

All data, code and models can be downloaded from https://github.com/Christini-Lab/iPSC-leak-artifact.

### 4.1 Modelled concentrations

I_leak_ in the baseline Kernik model destabilises intracellular concentrations and causes a slow and continuous decrease in [K^+^]_i_. To address this, we fixed the Kernik [K^+^]_i_ to its steady state value. This was not required for the Paci model, which already did not allow [K^+^]_i_ to change. We also fixed the Kernik and Paci [Na^+^]_i_ to their baseline steady state values (taken after 1000 s of spontaneous current clamp simulation using the published model initial values as starting points). We did not fix the intracellular calcium concentration, because it is less affected by the pipette solution during perforated patch-clamp experiments with amphotericin B as only monovalent ions can diffuse through the pores.

### 4.2 Genetic algorithm

A GA was used to fit the Kernik model to the Kernik+leak model (with R_seal_=5 GΩ) by minimising a point-by-point squared difference objective function:

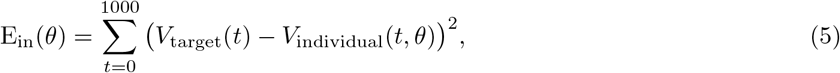

where *θ* is a vector containing the varied conductance parameters, *V*_target_(*t*) is the target membrane potential at time *t*, and *V*_individual_(*t, θ*) is the current individual’s membrane potential at time *t* as a function of *θ*.

The GA had a population size of 150 individuals and was run for 20 generations. The initial population parameter values were selected from a log uniform distribution between 0.1 and 10 times their baseline values. Individuals in a new generation were created by mating two individuals from the previous generation — two selected parent individuals from the previous generation had a 90% chance of mating. If they did not mate, they would continue to the next generation without swapping parameters. If they mated, there was a 20% chance of swapping each of their parameter values. As such, each time two individuals mated, they would produce two child individuals consisting of the parent parameter values. Each individual in a new generation had a 90% chance of being mutated. If an individual was mutated, there was a 20% chance each parameter would be changed. To mutate a parameter, a new value was selected from a normal distribution centred around the current value, with a standard deviation equal to 10% of the current value.

The Kernik+leak target and each individual were run for 100 s before comparison. The third-to-last AP was identified from each, and traces were aligned by the dV/dtmax of these APs. Traces were compared from 200 ms before the dV/dt_max_ to 800 ms after it. The code for this GA can be found on the project GitHub page.

### 4.3 Linear regression

A linear least-squares regression was used to compare *g*_in_/C_m_ to AP biomarkers. MP was the only biomarker that did not violate any linear regression assumptions when compared to these independent variables. Tests of these assumptions can be found on the project GitHub repository.

### 4.4 Software and simulations

Simulations were performed in Myokit v1.33.7 (Clerx et al., 2016). The genetic algorithm was developed in Python and made use of the DEAP library v1.2.2 (Fortin et al., 2012). Additional analysis was done in Python using NumPy v1.21.6 and SciPy v1.7.3 (Virtanen et al., 2020).

### 4.5 iPSC-CM cell culture

Frozen vials of iPSC-CMs were obtained from Joseph C. Wu, MD, PhD at the Stanford Cardiovascular Institute Biobank. The iPSC-CM line was derived from an African American female donor and the differentiation was approved by the Stanford University Human Subjects Research Institutional Review Board. Cells were prepared for electrophysiological experiments following the steps described in Clark et al. (2022). Briefly, cells were thawed and cultured as a monolayer in one well of a 6-well plate precoated with 1% Matrigel. Cells were cultured with RPMI media (Fisher/Corning 10-040-CM) containing 5% FBS and 2% B27 and kept in an incubator at 37 °C, 5% CO_2_, and 85% humidity. After 48 hours, cells were lifted with 1mL Accutase, diluted to 100, 000 cells/mL, and replated on 124 sterile 8 mm coverslips precoated with 1% Matrigel. Cells were cultured with RPMI media that was swapped every 48 hours. Cells were patched between days 5 and 15 after thaw.

### 4.6 Electrophysiological setup

Perforated patch-clamp experiments were conducted following the protocol described in Clark et al. (2022). Borosilicate glass pipettes were pulled to a resistance of 2-4 MΩ using a flaming/brown micropipette puller (Model P-1000; Sutter Instrument, Novato, CA). Pipette tips were first dipped into intracellular solution containing 10mM NaCl, 130mM KCl, 1 mM MgCl_2_, 10mM CaCl_2_, 5.5mM dextrose, 10mM HEPES. Pipettes were then backfilled with intracellular solution with 0.44 mM amphotericin B, a perforating agent. Amphotericin B allows only monovalent ions to pass through the cell membrane, so a high intrapipette calcium concentration was included to induce cell death in the case of an unintended rupture. Coverslips containing iPSC-CMs were placed in the bath and constantly perfused with an extracellular solution at 35-37°C containing 137 mM NaCl, 5.4 mM KCl, 1mM MgSO_4_, 2mM CaCl_2_, 10mM dextrose, and 10mM HEPES.

Patch-clamp measurements were made at a 10 kHz sampling frequency by an amplifier with the low-pass filter set to 5 kHz (Model 2400; A-M Systems, Sequim, WA), and was controlled by the Real Time eXperiment Interface (RTXI; http://rtxi.org). After immersing a pipette into the extracellular solution, voltage was set to zero — any remaining offset in the recordings is assumed to be equal to the liquid junction potential of −2.8 mV. After contact was made with a cell and a seal of > 300 MΩ was formed, the perforating agent slowly decreased the access resistance to the cell (usually 10–15 minutes). A series resistance of 9–50MΩ was maintained for all experiments. After gaining access, R_m_ at 0 mV was measured before and after acquiring AP data.

## References

Ahrens-Nicklas RC & Christini DJ (2009). Anthropomorphizing the mouse cardiac action potential via a novel dynamic clamp method. Biophysical Journal 97, 2684–2692.

Becker N, Horváth A, De Boer T, Fabbri A, Grad C, Fertig N, George M & Obergrussberger A (2020). Automated Dynamic Clamp for Simulation of IK1 in Human Induced Pluripotent Stem Cell–Derived Cardiomyocytes in Real Time Using Patchliner Dynamite8. Current Protocols in Pharmacology 88, 1–23.

Blinova K, Schocken D, Patel D, Daluwatte C, Vicente J, Wu JC & Strauss DG (2019). Clinical Trial in a Dish: Personalized Stem Cell–Derived Cardiomyocyte Assay Compared With Clinical Trial Results for Two QT-Prolonging Drugs. Clinical and Translational Science 12, 687–697.

Boyett MR & Fedida D (1988). A computer simulation of the effect of heart rate on ion concentrations in the heart. Journal of Theoretical Biology 132, 15–27.

Clark AP, Wei S, Kalola D, Krogh-Madsen T & Christini DJ (2022). An in silico-in vitro pipeline for drug cardiotoxicity screening identifies ionic proarrhythmia mechanisms. British journal of pharmacology 121, 303a–304a.

Clerx M, Collins P, de Lange E & Volders PGA (2016). Myokit: A simple interface to cardiac cellular electrophysiology. Progress in biophysics and molecular biology 120, 100–14.

Clerx M, Mirams GR, Rogers AJ, Narayan SM & Giles WR (2021). Immediate and Delayed Response of Simulated Human Atrial Myocytes to Clinically-Relevant Hypokalemia. Frontiers in Physiology 12, 1–21.

Doss MX, Di Diego JM, Goodrow RJ, Wu Y, Cordeiro JM, Nesterenko VV, Barajas-Martínez H, Hu D, Urrutia J, Desai M, Treat JA, Sachinidis A & Antzelevitch C (2012). Maximum diastolic potential of human induced pluripotent stem cell-derived cardiomyocytes depends critically on IKr. PLoS ONE 7.

Ebihara L & Johnson EA (1980). Fast sodium current in cardiac muscle. A quantitative description. Biophysical Journal 32, 779–790.

Fabbri A, Prins A & de Boer TP (2020). Assessment of the effects of online linear leak current compensation at different pacing frequencies in a dynamic action potential clamp system In 2020 Computing in Cardiology, pp. 1–4.

Feyen DA, McKeithan WL, Bruyneel AA, Spiering S, Hörmann L, Ulmer B, Zhang H, Briganti F, Schweizer M, Hegyi B, Liao Z, Pölönen RP, Ginsburg KS, Lam CK, Serrano R, Wahlquist C, Kreymerman A, Vu M, Amatya PL, Behrens CS, Ranjbarvaziri S, Maas RG, Greenhaw M, Bernstein D, Wu JC, Bers DM, Eschenhagen T, Metallo CM & Mercola M (2020). Metabolic Maturation Media Improve Physiological Function of Human iPSC-Derived Cardiomyocytes. Cell Reports 32.

Fortin FA, De Rainville FM, Gardner MA, Parizeau M & Gagné C (2012). {DEAP}: Evolutionary Algorithms Made Easy. Journal of Machine Learning Research 13, 2171–2175.

Goversen B, van der Heyden MAG, van Veen TAB & de Boer TP (2018a). The immature electrophysiological phenotype of iPSC-CMs still hampers in vitro drug screening: Special focus on IK1. Pharmacol Ther 183, 127–136.

Goversen B, Becker N, Stoelzle-Feix S, Obergrussberger A, Vos MA, van Veen TA, Fertig N & de Boer TP (2018b). A hybrid model for safety pharmacology on an automated patch clamp platform: Using dynamic clamp to join iPSC-derived cardiomyocytes and simulations of Ik1 ion channels in real-time. Frontiers in Physiology 8, 1–10.

Han L, Li Y, Tchao J, Kaplan AD, Lin B, Li Y, Mich-Basso J, Lis A, Hassan N, London B, Bett GC, Tobita K, Rasmusson RL & Yang L (2014). Study familial hypertrophic cardiomyopathy using patient-specific induced pluripotent stem cells. Cardiovascular Research 104, 258–269.

HEKA Elektronik GmbH (2016). Patchmaster multi-channel data acquisition software reference manual. Data Base 3304, 1–148.

Herron TJ, Da Rocha AM, Campbell KF, Ponce-Balbuena D, Willis BC, Guerrero-Serna G, Liu Q, Klos M, Musa H, Zarzoso M, Bizy A, Furness J, Anumonwo J, Mironov S & Jalife J (2016). Extracellular matrix-mediated maturation of human pluripotent stem cell-derived cardiac monolayer structure and electrophysiological function. Circulation: Arrhythmia and Electrophysiology 9, 1–12.

Horváth A, Lemoine MD, Löser A, Mannhardt I, Flenner F, Uzun AU, Neuber C, Breckwoldt K, Hansen A, Girdauskas E, Reichenspurner H, Willems S, Jost N, Wettwer E, Eschenhagen T & Christ T (2018). Low Resting Membrane Potential and Low Inward Rectifier Potassium Currents Are Not Inherent Features of hiPSC-Derived Cardiomyocytes. Stem Cell Reports 10, 822–833.

Jæger KH, Wall S & Tveito A (2021). Computational prediction of drug response in short QT syndrome type 1 based on measurements of compound effect in stem cell-derived cardiomyocytes. PLoS computational biology 17, e1008089.

Jonsson MK, Vos MA, Mirams GR, Duker G, Sartipy P, De Boer TP & Van Veen TA (2012). Application of human stem cell-derived cardiomyocytes in safety pharmacology requires caution beyond herg. Journal of molecular and cellular cardiology 52, 998–1008.

Kernik DC, Morotti S, Wu HD, Garg P, Duff HJ, Kurokawa J, Jalife J, Wu JC, Grandi E & Clancy CE (2019). A computational model of induced pluripotent stem-cell derived cardiomyocytes incorporating experimental variability from multiple data sources. Journal of Physiology 597, 4533–4564.

Koivumäki JT, Naumenko N, Tuomainen T, Takalo J, Oksanen M, Puttonen KA, Lehtonen Š, Kuusisto J, Laakso M, Koistinaho J & Tavi P (2018). Structural immaturity of human iPSC-derived cardiomyocytes: In silico investigation of effects on function and disease modeling. Frontiers in Physiology 9, 1–17.

Krogh-Madsen T, Schaffer P, Skriver AD, Taylor LK, Pelzmann B, Koidl B & Guevara MR (2005). An ionic model for rhythmic activity in small clusters of embryonic chick ventricular cells. American Journal of Physiology - Heart and Circulatory Physiology 289, 398–413.

Lei CL, Clerx M, Whittaker DG, Gavaghan DJ, de Boer TP & Mirams GR (2020). Accounting for variability in ion current recordings using a mathematical model of artefacts in voltage-clamp experiments. Philosophical transactions. Series A, Mathematical, physical, and engineering sciences 378, 20190348.

Lei CL, Fabbri A, Whittaker DG, Clerx M, Windley MJ, Hill AP, Mirams GR & de Boer TP (2021). A nonlinear and time-dependent leak current in the presence of calcium fluoride patch-clamp seal enhancer [version 2; peer review: 4 approved]. Wellcome Open Research 5, 152.

Lei CL, Wang K, Clerx M, Johnstone RH, Hortigon-Vinagre MP, Zamora V, Allan A, Smith GL, Gavaghan DJ, Mirams GR & Polonchuk L (2017). Tailoring mathematical models to stem-cell derived cardiomyocyte lines can improve predictions of drug-induced changes to their electrophysiology. Frontiers in Physiology 8.

Li W, Luo X, Ulbricht Y & Guan K (2021). Blebbistatin protects iPSC-CMs from hypercontraction and facilitates automated patch-clamp based electrophysiological study. Stem Cell Research 56, 102565.

Luo CH & Rudy Y (1994). A dynamic model of the cardiac ventricular action potential: I. Simulations of ionic currents and concentration changes. Circulation Research 74, 1071–1096.

Ma J, Guo L, Fiene SJ, Anson BD, Thomson JA, Kamp TJ, Kolaja KL, Swanson BJ, January CT, Kl K, Bj S & Ct J (2011). High purity human-induced pluripotent stem cell-derived cardiomyocytes: electrophysiological properties of action potentials and ionic currents. Am J Physiol Heart Circ Physiol 301, 2006–2017.

Mathur A, Loskill P, Shao K, Huebsch N, Hong SG, Marcus SG, Marks N, Mandegar M, Conklin BR, Lee LP & Healy KE (2015). Human iPSC-based cardiac microphysiological system for drug screening applications. Scientific Reports 5, 1–7.

Meijer van Putten RM, Mengarelli I, Guan K, Zegers JG, van Ginneken AC, Verkerk AO & Wilders R (2015). Ion channelopathies in human induced pluripotent stem cell derived cardiomyocytes: a dynamic clamp study with virtual IK1. Front Physiol 6, 7.

Mirams GR, Pathmanathan P, Gray RA, Challenor P & Clayton RH (2016). White paper: Uncertainty and variability in computational and mathematical models of cardiac physiology. The Journal of Physiology 594, 6833–6847.

Montnach J, Lorenzini M, Lesage A, Simon I, Nicolas S, Moreau E, Marionneau C, Baró I, De Waard M & Loussouarn G (2021). Computer modeling of whole-cell voltage-clamp analyses to delineate guidelines for good practice of manual and automated patch-clamp. Scientific Reports 11, 1–16.

O’Hara T, Virág L, Varró A & Rudy Y (2011). Simulation of the undiseased human cardiac ventricular action potential: Model formulation and experimental validation. PLoS Computational Biology 7.

Ortega FA, Grandi E, Krogh-Madsen T & Christini DJ (2018). Applications of dynamic clamp to cardiac arrhythmia research: Role in drug target discovery and safety pharmacology testing. Frontiers in Physiology 8, 1–8.

Paci M, Hyttinen J, Aalto-Setälä K & Severi S (2013). Computational Models of Ventricular- and Atrial-Like Human Induced Pluripotent Stem Cell Derived Cardiomyocytes. Annals of Biomedical Engineering 41, 2334–2348.

Pieske B, Maier LS, Piacentino V, Weisser J, Hasenfuss G & Houser S (2002). Rate dependence of [Na+]i and contractility in nonfailing and failing human myocardium. Circulation 106, 447–453.

ten Tusscher KH, Noble D, Noble PJ & Panfilov AV (2004). A model for human ventricular tissue. American Journal of Physiology - Heart and Circulatory Physiology 286, 1573–1589.

Virtanen P, Gommers R, Oliphant TE, Haberland M, Reddy T, Cournapeau D, Burovski E, Peterson P, Weckesser W, Bright J, van der Walt SJ, Brett M, Wilson J, Millman KJ, Mayorov N, Nelson ARJ, Jones E, Kern R, Larson E, Carey CJ, Polat b, Feng Y, Moore EW, VanderPlas J, Laxalde D, Perktold J, Cimrman R, Henriksen I, Quintero EA, Harris CR, Archibald AM, Ribeiro AH, Pedregosa F, van Mulbregt P & SciPy 1.0 Contributors (2020). SciPy 1.0: Fundamental Algorithms for Scientific Computing in Python. Nature Methods 17, 261–272.

Whittaker DG, Clerx M, Lei CL, Christini DJ & Mirams GR (2020). Calibration of ionic and cellular cardiac electrophysiology models. Wiley Interdisciplinary Reviews: Systems Biology and Medicine 12, e1482.

